# Two senses, one rhythm: Pre- and post-stimulus neural states predict perceived audio-visual synchrony

**DOI:** 10.1101/2025.05.20.655148

**Authors:** Siddharth Talwar, Francesca M. Barbero, Markus J. van Ackeren, Nathan Weisz, Olivier Collignon

## Abstract

How does the brain integrate or segregate multiple sensory inputs? In our study, we used magnetoencephalography to comprehensively investigate local and network dynamics before and after the presentation of sequences of beeps and flashes that are sometimes perceived as synchronous or asynchronous. Frequency-tagging and spectral analysis revealed that under identical physical conditions, subjective perceptual fusion of multisensory signals elicits stronger evoked sensory responses in both visual and auditory cortices. Crucially, perceptual fusion is preceded by locally reduced pre-stimulus alpha activity, as well as enhanced functional connectivity between auditory and visual cortices within the pre-stimulus gamma activity. We also show that pre-stimulus neuronal states in the occipital cortex predict post-stimulus sensory responses to auditory sequences in the temporal cortex, but not the opposite. The current findings provide evidence that the perceptual fusion of multisensory events relate to a complex interplay between local and inter-sensory neuronal states before and after stimulus presentation.

**Significance Statement:** How does the brain blend information from multiple sensory systems into one holistic perception? We developed a novel frequency-tagging paradigm in Magnetoencephalography (MEG) to robustly estimate the sensory responses to a series of flashes and beeps in early sensory areas, and to demonstrate how these responses are shaped by ongoing fluctuations in local, and long-range network activity. By combining the steady-state method with spectral, and functional connectivity analysis of ongoing oscillations in source space, the current study provides novel insights into the relationship between internal brain states and the way we perceive multisensory events.

## 1. Introduction

Understanding how information from multiple senses are seamlessly merged into a single coherent percept is a major challenge for the field of cognitive neuroscience. It is argued that integration emerges already at the earliest stages of the cortical hierarchy, within areas once considered exclusively unisensory.

To resolve conflicting sensory information the brain prioritizes more reliable sensory channels (Ernst and Banks, 2002). When localizing objects in space, we typically rely more on the visual system (e.g., ventriloquist effect, rubber hand illusion: Alais and Burr, 2004b; Ehrsson et al., 2004; Charbonneau et al., 2013), while audition typically dominates the detection of temporal changes (e.g., auditory capture: Shipley, 1964; Shams et al., 2000). A popular experiment to unravel the interaction between vision and audition is the double-flash illusion (DFI). Participants frequently perceive multiple visual flashes when a single flash is accompanied by multiple temporally interspersed auditory beeps (Shams et al., 2000). The DFI is a classic example where perception of one sensory system (vision) is influenced by another (audition). It is currently believed that such a perceptual effect may emerge already at the earliest stages of the auditory and visual cortical hierarchies (Shams & Kim, 2010), challenging the idea of a strict parcellation of the brain into unisensory and multisensory areas (Ghazanfar and Schroeder, 2006; Kayser and Logothetis, 2007; Lakatos et al., 2007; Driver and Noesselt, 2008).

Sensory areas are not passive gateways to the cortex. Rather, the neuronal states in early sensory systems as well as their long-range communication profiles with the rest of the brain shape the perception and integration of sensory events (Ebrahimi et al., 2022; Kayser & Logothetis, 2007). Several studies have demonstrated that neuronal states in early sensory systems can predict the detection of a weak sensory stimulus, or whether information from different input channels are combined (Lange et al., 2013, Lange et al., 2014; Keil et al., 2014; Leske et al., 2015). For instance, reduced pre-stimulus alpha power in the early visual cortices increases the likelihood of perceiving the DFI (Lange and colleagues (2013). Both local changes in the gain of sensory areas and modulations in sensory regions mediated by long-range functional connections can lead to fluctuations in excitability, thereby shaping perception (Ruhnau et al., 2014). While both local and long-range neuronal dynamics have been linked to multisensory integration (Lange et al., 2011, 2013, 2014;), it is not well understood how the interplay of local and global neuronal dynamics modulate perception.

We used Magnetoencephalography (MEG) and a novel frequency-tagging approach, combined with spectral and functional connectivity analysis of ongoing oscillatory activity that allows us to isolate sensory processing of a single modality in the frequency domain. Participants saw and heard sequences of flashes and beeps at different presentation frequencies, and were asked if the two sequences are in synchrony. Based on previous research on audio-visual illusions such as the flicker flutter effect (Shipley, 1964; Berger et al., 2003; Recanzone, 2003), we predicted that during some asynchronous trials, participants will perceive beeps and flashes to be in synchrony. Using frequency-tagging, we then isolated auditory and visual responses. Finally, the response magnitude was linked to the pre-stimulus neuronal states in each of the two systems. If pre-stimulus neuronal states operate locally, we predict a negative relationship between pre-stimulus alpha power and post-stimulus sensitivity to the sensory event in the *same sensory modality*. In contrast, if local fluctuations in alpha power bias the system at a more global level, we expect that local alpha power fluctuations in one sensory modality will affect sensitivity to sensory events in the *other modality*, potentially supported by enhanced long-range connectivity between sensory modalities.

## 2. Materials and Methods

### 2.1 Participants

Twenty-two healthy volunteers (10 males, 24-31 years) with no known neurological deficits participated in this study. Participants gave written informed consent and received a small monetary compensation. The project was approved by the local Ethical Committee at University of Trento. Due to a technical error, the behavioral data of one participant was not recorded and could not be considered in the analyses where trials were segregated based on the participant’s behavior.

### 2.2 Experimental Design

Visual stimuli were delivered into a dimly lit magnetically shielded room using a Dell Alienware Aurora PC running Windows 7 (64 bit). Stimuli were generated using Psychophysics toolbox 3 (Brainard, 1997) on Matlab 7.9 (The MathWorks Inc., Natick, MA). Images were projected onto a translucent screen using a PROPixx DLP LED projector at a refresh rate of 120Hz, and a screen resolution of 1920×1080 pixels. Participants viewed the stimulus (3.44° of visual angle) at a distance from the screen of 100cm. Auditory stimuli were presented via stereo loudspeakers using Panphonics Sound Shower 2 amplifier at a sound level that was audible and comfortable for the participant.

The stimuli included a Gaussian red dot, presented in the center of the screen at two different visual flashing frequencies (slow: 5Hz and fast: 7.5Hz) and a pure tone (500 Hz carrier frequency) that was amplitude modulated at three different auditory beep frequencies (5Hz, 6.5Hz, and 7.5Hz), resulting in six different flash-beep combinations. These combinations consisted of two physically synchronous (S) conditions (5.0V-5.0A and 7.5V-7.5A), and four asynchronous (AS) conditions (5.0V-6.5A, 5.0V-7.5A, 7.5V-5.0A, 7.5V-6.5A). The asynchronous conditions were further subdivided into two conditions where the difference between frequencies was far (ASf: 5.0V-7.5A, 7.5V-5.0A) versus close (ASc: 5.0V-6.5A, 7.5V-6.5A). The subjective experience of synchrony between the two sensory streams was expected to be stronger during the ASc condition (Fig. 1A, top panel). The choice for these presentation frequencies was based on the constraints introduced by the refresh rate of the screen (120Hz) and a series of behavioral pilot experiments. All S and ASf conditions comprised 80 trials, while the number of trials was doubled (160 trials) in the ASc conditions, to ensure a sufficient trial number when sorting into trials where participants perceptually integrated or segregated the two sensory events. In addition, 80 Null trials were included in the experiment for a baseline comparison. The experiment was divided into 10 blocks of 72 pseudo-randomized trials, resulting in a total of 720 trials.

**Figure 1.**
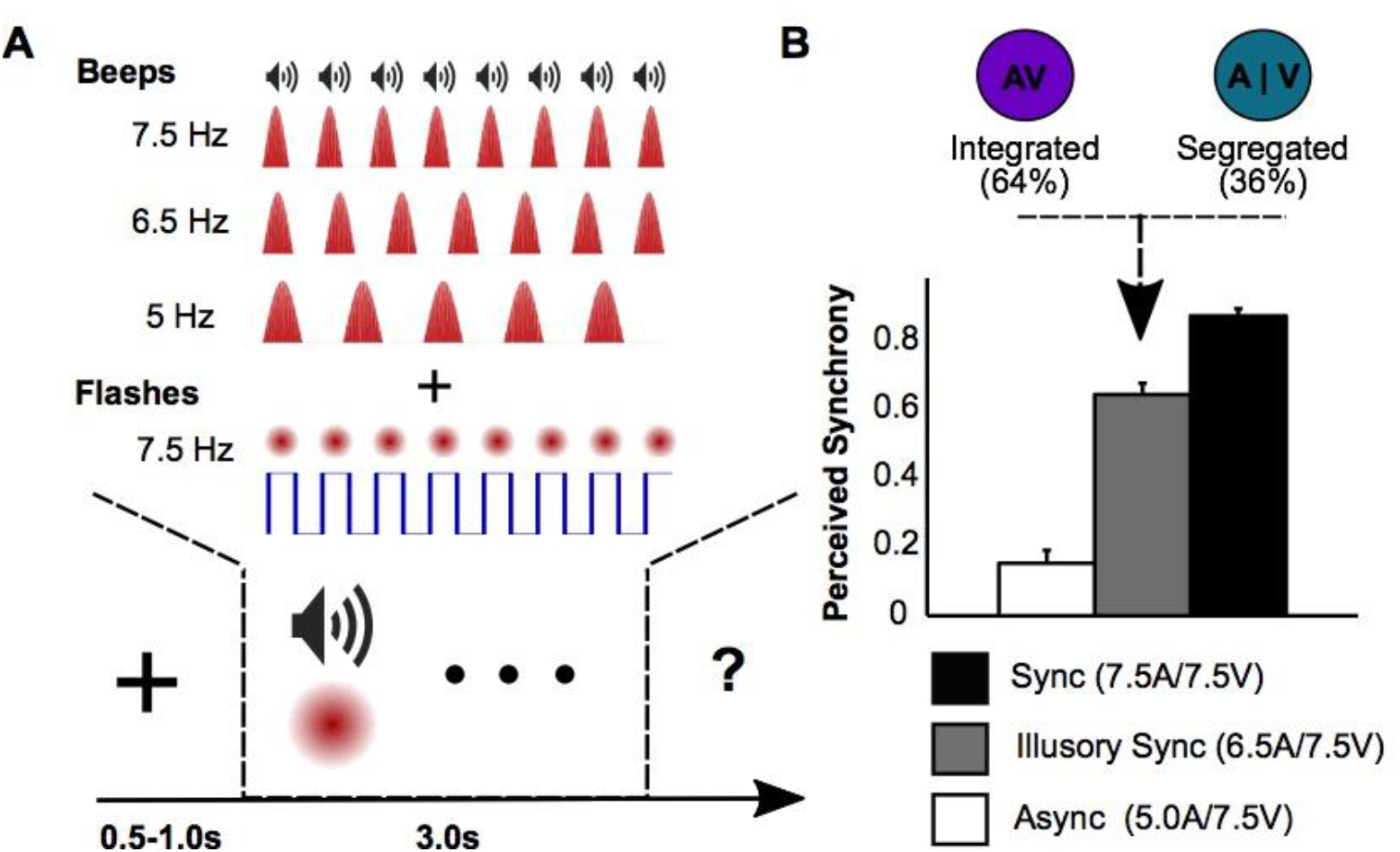
Design and behavioral results of the study. A) The top panel illustrates the set of conditions used in the current study. The rows depict the three beep frequencies (5Hz, 6.5Hz, 7.5Hz), while the columns show the two flash frequencies (5Hz, 7.5Hz). The bottom panel shows the temporal sequence of a single trial. After a variable period of fixation, auditory beeps and visual flashes are presented at variable frequencies for 3 seconds. A trial is always followed by the question if the two sensory sequences were in synchrony or not. B) The behavioral results for the fast visual presentation frequency (7.5 Hz) show a gradual increase in the proportion trials where the two events were perceived as synchronous as the beep frequencies approached the flash frequency. In the ASf condition, synchrony is perceived on 64% of the trials. Error bars depict SEM, adjusted for within subject comparisons.

Participants were given a two-alternative forced choice task (2AFC). After a period of fixation (750±250 ms), the audio-visual stimuli were presented for 3 sec. Participants were asked to indicate whether the beeps and flashes were in synchrony (Fig. 1A, bottom panel).

### 2.3 MEG Acquisition and Preprocessing

MEG was recorded continuously on a 102 triple sensor (two gradiometer, and one magnetometer) whole-head system (Elekta Neuromag, Helsinki Finland). Data was acquired with a sampling rate of 1 kHz and an online band pass filter between 0.1 and 330 Hz. The headshape of each individual participant was measured using a Polhemus FASTRAK 3D digitizer. The head position was recorded continuously using five localization coils (forehead, mastoids). For source reconstruction, individual structural MRI images of 12 participants were acquired on a 4T scanner (Bruker Biospin Ettlingen, Germany) The images were consequently co-registered into the MEG coordinate system using a set of anatomical landmarks (pre-auricular points and nasion) as well as the individual headshape. When no individual structural MRI was available, a model of the individual anatomy was created by warping an MNI template brain (MNI, Montreal, Quebec, Canada; http://www.bic.mni.mcgill.ca/brainweb) to the individual subject’s headshape (10 participants). The structural MRI was used to create a realistic single-shell headmodel (Nolte, 2003).

The data preprocessing and analysis was performed using the open-source toolbox fieldtrip (Oostenveld et al., 2011), as well as custom written code. The continuous data was filtered (high-pass Butterworth filter at 1Hz; DFT filter at 50, 100, and 150 Hz) and downsampled to 500 Hz. The resulting continuous data was epoched around the events of interest and inspected visually for ocular, muscle and jump artifacts. Subsequently, cleaned epochs were band-pass filtered, depending on the analyses to be performed, and projected onto a 3-dimensional grid (1.5×1.5×1.5cm) using a linearly constrained minimum variance (LCMV) beamformer (Van Veen et al., 1997).

### 2.4 Spectral analysis in source space

Analysis of steady-state responses in source space was performed on the band-pass filtered data (4-16Hz), and centered on a covariance window between 1000-3000 ms post-stimulus to avoid evoked activity in the signal during the first second. Individual trials in the time window of interest were baseline corrected, averaged in the time domain, and subjected to a Fast Fourier transformation (FFT). The resulting complex values were used to compute the amplitude of the different frequencies.

Time-resolved Fourier spectra of pre-stimulus oscillatory activity were computed in source space, separately for low (4-32Hz) and high frequencies (32-128Hz). For low frequencies, the data was band-pass filtered (4-32Hz) and projected into source space using a covariance window of 600 ms before stimulus onset. Spectral components were estimated in 0.25 octave steps using a single Hanning taper on 500ms sliding time windows (50ms step size). High frequency spectral components were estimated from the high-pass filtered signal (32Hz) in source space using a multitaper approach with three Slepian tapers and sliding time windows of 250ms (Mitra and Pesaran, 1999). This procedure resulted in a frequency resolution of ±12.5Hz. The same Fourier spectra were used to test for differences in pre-stimulus spectral power and connectivity. Functional connectivity was operationalized with the Phase Locking Value (PLV) (Lachaux et al., 1999), which quantifies the consistency in phase angle differences between two regions, independent of the amplitude.

### 2.5 Regression analysis

A linear regression approach was used to study the potential relationship between pre-stimulus alpha power fluctuations in early sensory areas, and post-stimulus activity at the auditory and visual tagging frequencies. First, single trial alpha power was computed at the source level in early visual and auditory areas. Next, trials were binned into quartiles on the basis of single-trial pre-stimulus alpha power, and post-stimulus steady state responses were computed across all trials in each of the four bins. Finally, linear regression was used at all grid points to find which brain regions show an inverse relationship between the alpha power in early auditory and visual areas and post-stimulus activity at the auditory and visual presentation frequencies. To control for multiple comparisons in space, a cluster-based resampling procedure was applied based on the cluster-mass. Correction for multiple comparisons was applied using a maximum statistic; the observed cluster-mass was compared to a permutation distribution of the maximum cluster-mass value from each permutation.

### 2.6 Statistical analysis

Statistical analysis of frequency-tagged responses was performed using permutation t-tests (1000 iterations). At the whole brain level, a multiple comparison correction was applied based on the false discovery rate (FDR, Benjamini and Hochberg, 1995; Genovese et al., 2002). For the analysis of pre-stimulus oscillatory activity, second-level random-effects analysis was conducted. First, independent-sample t-tests were computed between conditions for each participant. The resulting t-values were converted to z-values for second level analysis. These z-maps were tested against zero at the group level. To control for multiple comparisons in time-frequency space, a cluster-based permutation approach was used, based on a maximum statistic of the cluster-mass (Maris and Oostenveld, 2007). The same resampling procedure was used to correct for multiple comparisons in space for the pre-post regression analysis.

## 3. Results

### 3.1 Behavioral Performance

Paired-sample permutation t-test (1000 iteration) revealed that participants perceived beeps and flashes as synchronous significantly more often during the Synchronous (S) versus the Asynchronous Far (ASf) conditions for slow (5.0V-5.0A versus 5.0V-7.5A Hz: *t*(20)=8.13, *p*<.001) and fast visual presentation frequencies (7.5V-7.5A versus 7.5V-5.0A Hz: *t*(20)=14.00, *p*<.001). Furthermore, synchrony was reported more often during the Asynchronous Close (ASc) versus Asynchronous Far (ASf) condition for fast (7.5V-6.5A versus 7.5V-5.0A Hz: *t*(20)=7.28, *p*<.001), but not for slow visual presentation frequencies (5.0V-6.5A versus 5.0V-7.5A Hz: *t*(20)=0.34, *p*=.774). Taken together, the behavioral results show that we were able to reliably elicit the subjective illusion of synchrony in the ASc condition for fast (7.5Hz), but not for slow (5.0Hz) visual presentation frequencies (see Fig. 1b). Based on these results, the MEG analysis on the neuronal dynamics during subjective experience of synchrony targeted the fast visual presentation frequencies only (7.5 Hz).

### 3.2 Modality-Specific Responses to Physical and Perceived Audio-Visual Synchrony

To isolate early auditory and visual regions in space, frequency spectra of the ASf conditions (7.5V-5.0A and 5.0V-7.5A) were contrasted directly at the source level. For the localization, we used the 5Hz tagging frequency as it produced the strongest steady-state response in our data, independent of conditions (the results for 7.5Hz are slightly weaker but show a similar spatial distribution). As expected, we observe robust auditory 5Hz responses (7.5V-5.0A) in bilateral auditory cortex, while the visual 5Hz frequency (5.0V-7.5A) is strongest around V1 (Fig 2A). Regions of interest (ROI) for subsequent analysis were selected as the grid points in anatomically defined primary visual, and auditory cortex (www.brainmap.org), which were sensitive to the visual and auditory presentation frequencies respectively (FDR-corrected). This was done to reduce confounding effects from adjacent temporal and parietal regions, as the unmasked statistical maps also extended into association cortices.

**Figure 2.**
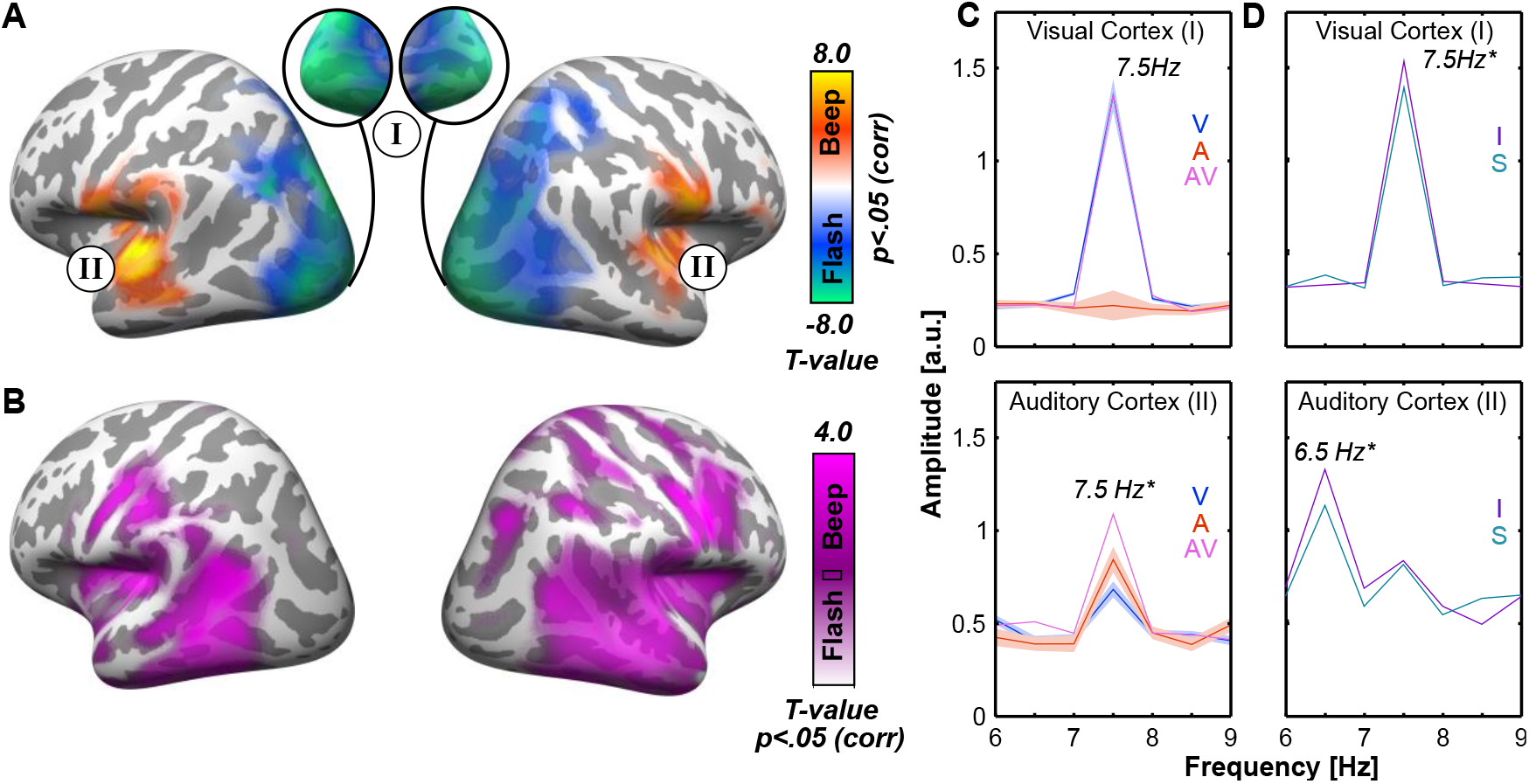
Analysis of auditory and visual processing at the presentation frequencies. A) The contrast between the two ASf conditions (7.5V–5.0A versus 5.0V-7.5A), at 5Hz was used to localize early sensory regions of interest. Warm colours show sensitivity to the rhythmic auditory stimulation in bilateral superior temporal gyrus. Cold colours reflect sensitivity to the visual presentation frequency in occipital and parietal lobes, with peaks in bilateral calcarine sulci. The statistical maps are thresholded at p<.05 (FDR-corrected). B) The activation maps show the multisensory areas from the conjunction between each of the two asynchronous against the Null condition. Minimum t-values are used to show gradient activation. C) Comparison of the S condition (7.5V-7.5A: pink) versus the two ASf conditions (7.5V-5.0A: blue 5.0V-7.5A red) in visual (top panel), and auditory (bottom panel) cortices. A synchrony effect is observed only in the auditory ROI. Shaded areas around the graphs depict SEM, adjusted for within subject comparisons. D) Illusory synchronization effects for the ASc condition during fast visual presentation (7.5V6.5A) in visual (top panel), and auditory (bottom panel) cortices. Enhanced neural activity is observed in both modalities for trials where visual and auditory signals were perceived as synchronous (purple) versus asynchronous (blue). Shaded areas around the graphs depict SEM, adjusted for within subject comparisons.

While the functional peaks of the frequency-tagged responses are located in primary sensory areas, they potentially may not be restricted to these regions. To explore this possibility, we contrasted the source amplitude at 5Hz for each ASf condition (5.0V-7.5A, max response in visual cortex; 7.5V-5.0A, max response in auditory cortex) against the Null-condition and computed a conjunction analysis (Fig. 2B). We observe a widespread overlap between the responses to auditory and visual signals in temporal and parietal areas that are classically considered multisensory integration zones (Beauchamp et al., 2004; Bremmer et al., 2004; Beauchamp, 2005).

The behavioral data revealed that the subjective illusion of synchrony was only reliably perceived in the ASc condition during fast (7.5V-6.5A) but not slow (5.0V-6.5A) visual presentation frequencies. Therefore, subsequent ROI-based analyses were based on the fast visual presentation frequencies (S: 7.5V-7.5A; ASc: 7.5V-6.5A) as well as the two ASf conditions (7.5V-5.0A, and 5.0V-7.5A).

For the ROI analysis, paired-sample permutation t-tests were computed at the presentation frequency (7.5Hz) to test whether physical synchrony of beeps and flashes alters neural responses in early sensory areas. In the auditory cortex, we observed a positive difference between the S (7.5V-7.5A) versus ASf (5.0V-7.5A) condition, suggesting that a stronger neuronal response when the two sensory streams are in synchrony (*t*(21)=4.23, *p* = .001; Fig. 2C). In contrast, there was no evidence for an effect of synchrony in the visual ROI (*t*(21)=0.69, *p*=.942). Taken together, we demonstrate that the auditory, but not visual cortex is sensitive to the physical synchrony between the two sensory streams.

Next, we tested whether auditory and visual cortices were sensitive to the subjective illusion of synchrony in the ASc condition (7.5V-6.5A), the AS condition which yielded the highest number of behavior responses indicating synchrony. On average, participants perceived the two events as synchronous on 64% of the trials (see Fig. 1B). Prior to statistical inference, an equal number of trials were randomly selected from the ASc condition where participants reported the two events to be synchronous (integrated) versus asynchronous (segregated). This was done to avoid any spurious effects due to different sample sizes. To enhance robustness of the tagged frequency amplitudes, participants were only considered for this analysis if more than 20% of the possible number of trials in a condition could be retained. This procedure resulted in the trimming of the data where participants responding predominantly with one type of response (either synchronous or asynchronous) were excluded (5 participants).

Paired-sample permutation t-tests were computed at the presentation frequencies (visual: 7.5Hz; auditory: 6.5Hz) on the remaining 16 participants. For the contrast between integrated versus segregated trials, we observed a significant difference in the auditory cortex (6.5Hz; *t*(15)=2.49, *p*=.021) (see Fig 2C). Specifically, the auditory cortex responds more strongly if the two sensory streams are integrated and perceived as synchronous. The same contrast at the visual tagging frequency (7.5Hz) demonstrated an equivalent effect in the visual ROI (*t*(15)=2.57, *p*=.019) (Fig 2D). Thus, the visual cortex responds more strongly to visual information when it is subjectively integrated with the auditory stream.

Taken together, the auditory cortex showed enhanced sensory responses to auditory information if the two sensory streams were either physically or perceptually in synchrony. In contrast, the visual cortex showed enhanced sensory responses only if the two asynchronous streams were perceived as synchronous, but not when the stimuli were physically in synchrony.

### 3.3 Pre-Stimulus Neuronal States Predict Multisensory Binding

Next, we tested whether neuronal states of early sensory areas preceding stimulus presentation predict whether auditory and visual streams are perceptually integrated during the ASc condition (7.5V-6.5A, the AS condition which yielded the highest number of behavior responses indicating synchrony). To this end, we performed a second-level random-effects analysis. First, independent-samples t-tests were computed for each individual subject contrasting *integrated versus segregated* trials, and the resulting t-values were converted to z-values for group-level analysis. At the group level, these z-maps were tested against zero using one-sample t-tests. Cluster-based permutation was performed to control for multiple comparisons in time and frequency space (Maris and Oostenveld, 2007). Analyses were conducted on visual and auditory ROI’s, separately for low (4-32Hz) and high frequencies (38-128Hz) in a time window of 500-100 ms before stimulus onset.

In the visual cortex (Fig 3A), we observed a reduction in spectral power around the alpha band (∼11.4 Hz) when participants perceived the sensory events as synchronous (*t*_*sum*_(15)=-91.61, *p*=.002). Furthermore, we found enhanced visual gamma activity (∼64-90.6Hz) in the same region when participants reported seeing the two sensory streams as synchronous (*t*_*sum*_(15)=4.24, *p*=.042). Thus, reduced alpha and enhanced gamma power in the visual cortex predict the perception of synchrony between auditory and visual events in the ASc condition.

**Figure 3.**
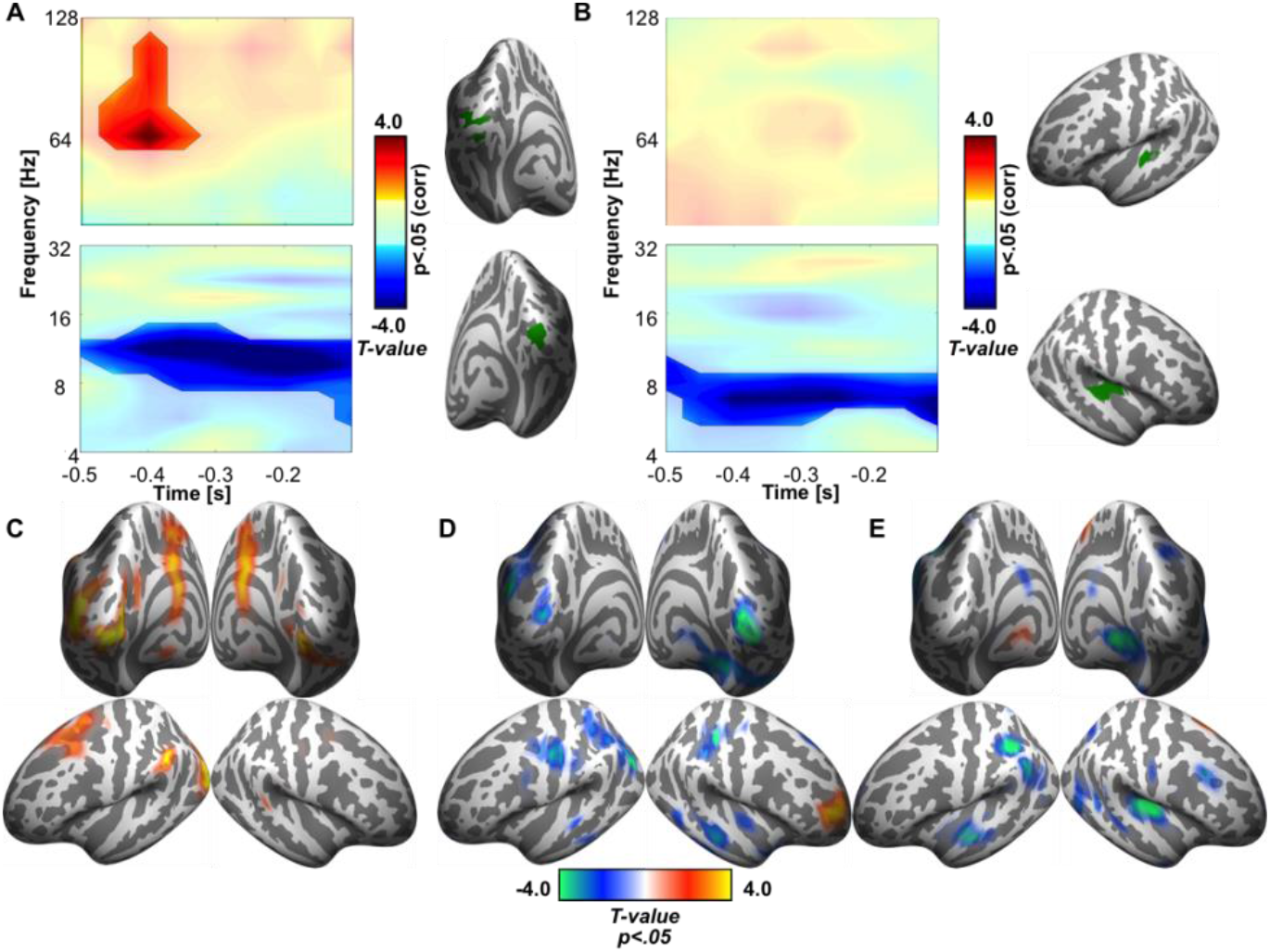
Analysis of pre-stimulus oscillatory activity contrasting trials perceived as synchronous versus asynchronous. Time-frequency representations from visual areas (right panel) depict the statistical differences in pre-stimulus power for the integrated versus segregated trials. A) statistically significant increase (highlighted, top panel) is observed in the gamma band when audio-visual streams were integrated. In addition, there was a reduction in pre-stimulus alpha power B)) In auditory areas (right panel), there was a reduction in pre-stimulus alpha activity when the two sensory streams were perceived as synchronous (highlighted, bottom panel). C-E Whole-brain, statistically thresholded (p<.05, unc.) maps of the three effects in the visual gamma (C), visual alpha (D), and auditory alpha (E) band reveal mostly independent clusters in visual and auditory areas respectively. Additional clusters are observed mostly in temporal and parietal areas, however the peaks are found in the primary sensory regions.

In the auditory cortex (Fig. 3B), our analysis revealed a reduction in spectral power in the lower alpha band (∼6.8Hz) when trials were reported as synchronous (*t*_*sum*_(15)=-72.90, *p*=.006). Notably, the peak difference was at a lower frequency than what we observed in the visual cortex. Although there is some variability in the reported auditory alpha peak frequency (Weisz et al., 2011; Weisz and Obleser, 2014), some studies suggest that it is lower than the visual alpha rhythm (Lehtelä et al., 1997). In contrast to the visual cortex, no differences were found in the higher frequencies when participants perceived the trials as synchronous or asynchronous. Taken together, we observed that reduced auditory alpha power predicts the subjective experience of audio-visual synchrony in the auditory cortex.

To explore the spatial specificity of these effects, pre-stimulus power in the two conditions were computed for all points in the 3D grid whereby spectral power was averaged in the time-frequency window where the effect was maximal in the ROI analysis (visual alpha: 9.6-11.4Hz, 500-250ms; auditory alpha: 6.8-8Hz, 500-250ms; visual gamma: 64-90.6Hz, 500-350ms). The thresholded maps for the contrast between integrated versus segregated trials revealed peak differences in visual cortices for visual gamma and visual alpha and a peak in the right auditory cortex for auditory alpha (Figure 3C-E, respectively). The majority of additional peaks outside early sensory areas were located in parietal and temporal areas, largely overlapping with the audio-visual conjunction map.

Finally, to test whether there is a relationship between pre-stimulus neuronal states and post-stimulus evoked responses to the stimulus, we used a regression-based approach on the pre-stimulus alpha power, and post stimulus evoked activity at the presentation frequencies in early visual and auditory cortex. First, alpha power was averaged across the time window 500-250 ms before stimulus onset. The decision for this time window was chosen a) to capture the maximum alpha suppression effect from the ROI analysis, and b) to avoid potential leakage from the post-stimulus time window into the pre-stimulus period due to the temporal smoothing of the time-frequency analysis (with a 500ms time window, post-stimulus events may leak into the pre-stimulus period up to 250ms). Averaged pre-stimulus alpha power in early sensory regions was computed for each individual trial, and trials were binned into quartiles. For each of these quartiles we also computed the amplitude at the auditory and visual presentation frequency post-stimulus. This was done for all points in the 3D grid. The procedure was applied separately with bilateral visual, and auditory cortices as seed regions, taking into account the preferred alpha rhythm in each modality (visual 9.6-11.4 Hz; auditory 6.8-8Hz). Statistical analyses were performed using linear regression, and cluster-based permutation was used to control for multiple comparisons (Maris and Oostenveld, 2007; Obleser and Weisz, 2012). Importantly, amplitudes of the presentation frequencies could show a linear increase or decrease with pre-stimulus alpha activity. However, the interpretation of an increase would be highly problematic as it could merely reflect noise fluctuations from trial to trial. Therefore, statistical analysis was performed one-sided to identify negative linear relationships only.

Our analysis revealed a significant negative relationship between the pre-stimulus visual alpha rhythm and the amplitude of the visual tagging frequency (7.5Hz: *t*_*sum*_ (21)=-165.33, *p*=.009). This difference was observed in posterior parietal cortex, predominantly in the right hemisphere, as well as in parieto-occipital regions stretching from extrastriate visual cortex medially towards the Precuneus and cingulate cortex (Fig. 4A,C). Furthermore, we also find a negative linear relationship between pre-stimulus visual alpha power and the auditory presentation frequency (*t*_*sum*_ (21)=-107.38, p=.026). This effect was observed in the right auditory cortex (Fig. 4B,D). There was no evidence for a negative linear relationship between the auditory alpha rhythm on either visual or auditory tagging frequency. Taken together, these results demonstrate that reduced visual alpha power in the early visual cortex predicts enhanced sensitivity to the flash sequence in downstream parieto-occipital areas, as well as enhanced auditory cortex sensitivity to the beep sequence. As no such effect was observed for the auditory alpha rhythm, we can conclude that in the case of auditory capture, there is an asymmetric relationship between the effects of locally reduced alpha power and cortical excitability in visual and auditory cortices.

**Figure 4.**
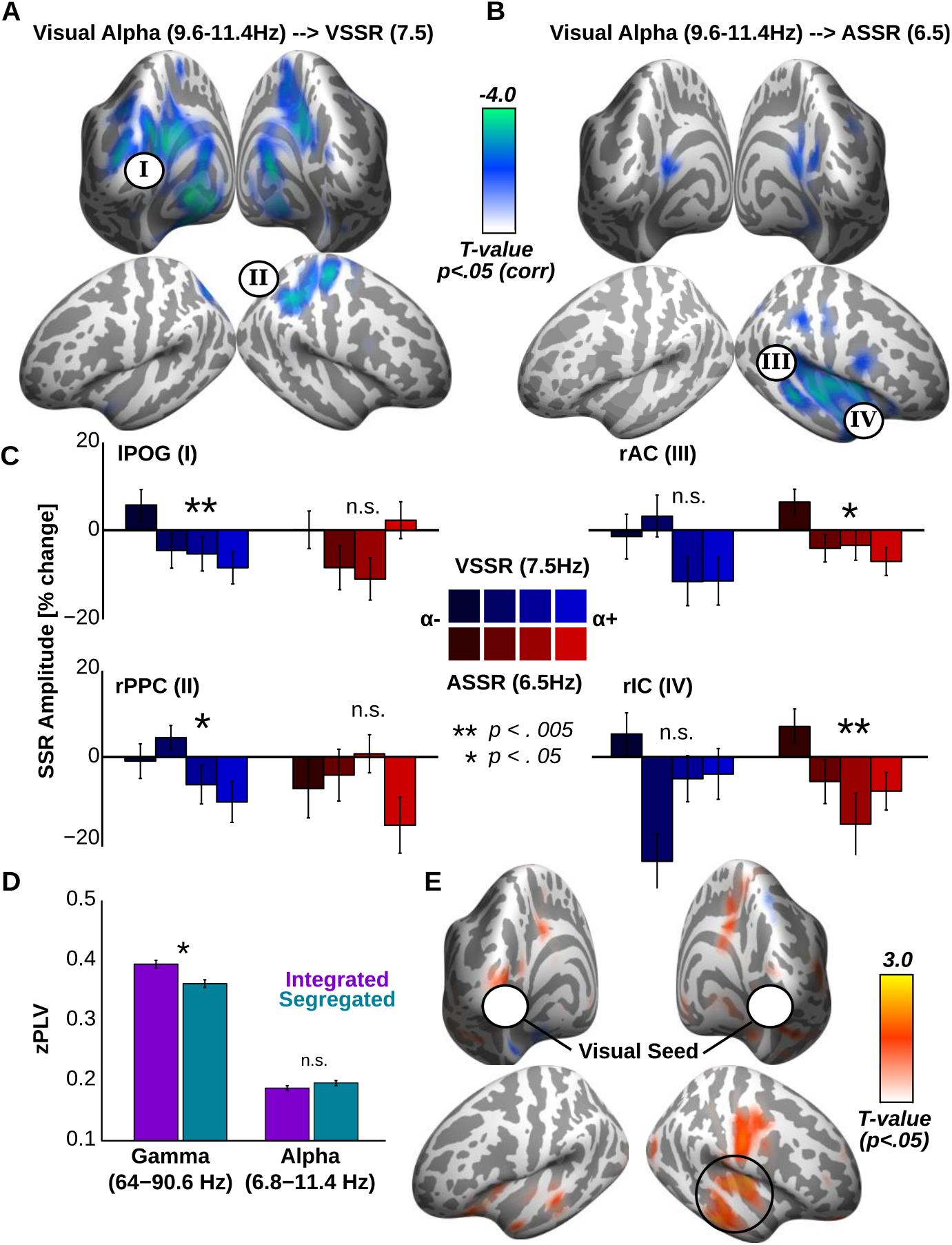
Pre-post regression and connectivity analysis. A) Statistically thresholded whole-brain activation map (p<.05, cluster-corrected, one-tailed), showing regions where the post-stimulus amplitude at the visual tagging frequency shows a negative linear relationship with pre-stimulus alpha power in visual cortex. This is observed in a network of regions spanning from parieto-occipital regions (POG: I) to posterior parietal cortex (rPPC: II) B) Activation maps show regions where a linear negative relationship is observed between the amplitude at the auditory tagging frequency post-stimulus, and pre-stimulus alpha power in visual cortex. This relationship is observed most strongly in the right auditory cortex (rAC: III) and insular cortex (rIC: IV). C) Bar graphs illustrate mean post-stimulus amplitude for the visual (blue) and auditory (red) tagging frequencies in each of the alpha quartiles from visual cortex (low alpha: dark shades, high alpha: lighter shades). Values are extracted from two peak regions where alpha is linearly related to the visual tagging frequency (lPOG, rPPC, left panel), and two regions where alpha is linearly related to the auditory tagging frequency (rAC, rIC, right panel). A significant negative linear relationship is marked with a star. Error bars depict SEM adjusted for within subject comparisons. D) Bar graphs show phase-locking between visual and right auditory cortex in the alpha and gamma band for integrated versus segregated trials. Enhanced functional connectivity is observed for integrated trials in the gamma, but not the alpha band. E) Thresholded statistical maps (p<.05, unc.) contrasting functional connectivity maps between the integrated versus segregated conditions considering the visual cortex as the seed region. The map shows that the activity is largely confined to the right superior temporal gyrus and other auditory regions.

If pre-stimulus alpha power in visual cortex predicts post-stimulus auditory processing, there might be pre-existing fluctuations in how the visual and auditory cortices communicate with each other that influence whether the two sensory events are perceived as synchronous (Ruhnau et al., 2014; Weisz et al., 2014; Leske et al., 2015). To directly test this idea, we computed the phase locking value (PLV), as a proxy for functional connectivity, between the right auditory and visual ROI for the visual alpha (6.8-11.4Hz, 500-250ms) and gamma band (64-90.6Hz, 500-250ms). Integrated and segregated trials were contrasted using paired-sample permutation t-tests. We found enhanced pre-stimulus functional connectivity between visual and auditory cortices in the gamma band on trials that were perceived as synchronous (*t*(15)=3.16, *p*=.006). To explore the spatial selectivity of this effect, permutation t-tests were computed contrasting the integrated versus segregated conditions at the whole-brain level. As the thresholded map (*p*<.05, unc.) in Fig. 4E illustrates, the strongest and most widespread increase in connectivity was observed between visual cortex and right auditory cortex, largely overlapping with the cluster from the regression analysis. Paired-sample t-tests in the alpha range did not yield a significant difference between integrated versus segregated trials (*t*(15)=0.66 *p*>.5) (Fig. 4D).

## 4. Discussion

We aimed to investigate the network mechanisms underlying the integration of audiovisual sequences by quantifying local representations and connectivity of early auditory and visual cortices. To this end we used a novel frequency tagging approach inspired by auditory capture effects such as the flicker flutter illusion (Ogilvie, 1956; Shipley, 1964; Berger et al., 2003; Noesselt et al., 2008); where a series of beeps and flashes are presented at neighboring frequencies, and the two streams are sometimes perceived as synchronous. Leveraging the intra-individual variability in multisensory perception, we studied physiological responses within and between auditory and visual systems in different perceptual contexts, that is, when the events were perceived as synchronous or not. Moreover, by linking these isolated sensory responses with ongoing oscillatory activity before the stimulus was presented, we were able to investigate how the neuronal state of one system affects the local sensory response, as well as the sensory response in the other sensory system. Such a paradigm and analytical framework allowed us to isolate the brain correlate of multisensory integration versus segregation under constant physical stimulation.

First, we observed that the auditory cortex responded more strongly to the auditory stream when the two sensory events were physically synchronous (Fig 2C, bottom panel), which was not true for the visual cortex. We also found enhanced auditory responses when participants perceived an asynchronous series of beeps and flashes as synchronous (Fig 2D, bottom panel). A similar enhancement in the visual response to the flashes emerged when participants report perceiving a stream of asynchronous beeps and flashes as synchronous (Fig 2D, top panel). Furthermore, the perception of synchrony is predicted by fluctuations in oscillatory neuronal activity shortly before stimulus presentation (Fig 3). In the auditory cortex pre-stimulus alpha power is reduced when participants integrate the two sensory events (Fig 3B,E). In the visual cortex, both reduced alpha and enhanced gamma power predict perceptual integration (Fig A,C,D). Most importantly, we demonstrated that these pre-stimulus dynamics have a different relationship with post-stimulus processing in visual and auditory cortices (Fig 4). Reduced visual alpha power predicts enhanced sensory processing in both auditory and visual cortices (Fig 4A-C), while we found no evidence for a relationship between pre-stimulus alpha power and post-stimulus sensory processing in the auditory cortex. Corroborating this idea of direct information transfer between early sensory areas, we show that the subjective perception of synchrony is preceded by enhanced functional connectivity in between sensory areas in the gamma band (Fig 4D-E).

### Sensory Processing During Physical and Perceptual Synchrony of Audio-Visual Events

In the current study, the auditory cortex responded more strongly if auditory and visual events were either physically or perceptually synchronous (Fig 2C-D, bottom panel). In contrast, the visual cortex showed enhanced sensory responses only when the two sensory streams were perceptually synchronous and not physically synchronous (Fig 2D, top panel). This pattern of results is interesting because it attributes different roles to auditory and visual cortices with respect to multisensory integration: While auditory cortex appears to be sensitive to both perceptual and physical synchrony, the response in visual cortex is only linked to the illusory perception of synchrony. These results may relate to the auditory capture, as ‘integration’ of the two sensory streams shift in perception toward the more reliable auditory stream (Recanzone, 2003). It is important to emphasize that although this asymmetry might hold for auditory capture, it is not likely generalizable to other forms of multisensory integration.

Previous studies on synchrony effects in multisensory paradigms using frequency tagging are rare and have produced mixed results (Giani et al., 2012; Nozaradan et al., 2012; Barbero et al., 2025). Nozaradan and colleagues (2012) reported synchrony effects for both auditory and visual presentation frequencies. Moreover, intermodulation frequencies (linear combinations of audio-visual presentation frequencies) were proven to be a marker of audio-visual emotion integration (Barbero et al., 2025). As these results are restricted to sensor space it is not clear whether the effects reside in early sensory areas or in regions in association cortices. Indeed, using conjunction analysis we found a wide network of areas that are sensitive to both the visual and auditory presentation frequencies (Fig. 2B), which could result in multisensory effects observed at the sensor level. Our approach is different as it uses source analysis in a way that preserves the ability to maximally segregate the two sensory events using frequency tagging. By applying this more sensitive approach, we demonstrated that frequency tagging is a versatile tool to target multisensory integration in the brain.

### Excitability Fluctuations in Early Sensory Regions Link to Local and Network Activity Change

The current study further demonstrates that illusory perception of synchrony is associated with reduced power in the pre-stimulus alpha rhythm in both auditory and visual cortices (Fig 3). Current theories suggest that fluctuations in alpha power reflect changes in cortical excitability (Romei et al., 2008a, 2008b, 2010; Lange et al., 2013; Cecere et al., 2014). In the visual cortex, reduced alpha power is accompanied by enhanced visual gamma activity, often associated with visual processing (Gray et al., 1989; Womelsdorf et al., 2007; Fries et al., 2008). We demonstrated a negative relationship between pre-stimulus alpha power in visual areas and post-stimulus processing at the visual presentation frequency in downstream visual areas in parieto-occipital cortex (Fig 4A,C, left panel), as well as with the post-stimulus processing at the auditory presentation frequency in auditory regions (Fig 4B,C, right panel). Importantly, the reduced pre-stimulus alpha power in the occipital cortex predicted subjective multisensory integration (Fig. 3A,B). The current findings show that fluctuations in alpha power not only reflect local changes in cortical excitability, but also shape the dynamics of transient functional networks at large.

During the double flash illusion (Shams & Kim, 2010), it was argued previously that a second beep elicits the illusory perception of an additional flash when excitability in the visual cortex is high (Lange et al., 2013, Lange et al., 2014). In this study, the perceived synchrony could be predicted by the pre-stimulus alpha, which signifies cortical excitability. Furthermore, only the visual, but not the auditory alpha rhythm could be linked reliably to the amplitude of the tagged visual frequency, suggesting that in the case of auditory capture alpha power in the visual cortex has more direct effects on the global network organization, while auditory alpha operates locally.

Another interpretation of the excitability hypothesis is that reduced local alpha power makes visual and auditory cortices more prone to communicate (Lange et al., 2014). Two findings in the current study provide support for this idea. First, we found that enhanced functional connectivity in the gamma frequency band between early visual and auditory areas predicted whether the two sensory events are perceived as synchronous (Fig 4D-E). This is in line with animal studies demonstrating the existence of direct anatomical pathways between early auditory and visual areas (Falchier et al., 2002; Rockland and Ojima, 2003; Cappe and Barone, 2005). Second, and most importantly, we observed that reduced pre-stimulus alpha power in the visual cortex predicts post-stimulus sensory responses in both visual and auditory cortices (Fig 4B-C, right panel). In contrast, local alpha fluctuations in the auditory cortex, while linked to the perception of synchrony, do not affect the sensory response in the visual cortex. This asymmetric relationship between reduced pre-stimulus alpha power and enhanced post-stimulus sensory responses in visual and auditory cortices could arise from the fact that vision is our dominant modality and under most circumstances drives multisensory perception. As such spontaneous fluctuations in the visual cortex will affect both local and global network structure more readily than fluctuations in the auditory cortex.

In conclusion, our study demonstrates that temporal integration of multisensory events in time depends on the fluctuating neuronal states of the system, which bias perception in early sensory areas through local and long-range network interactions.

